# Future flooding tolerant rice germplasm: resilience afforded beyond *Sub1A* gene

**DOI:** 10.1101/2025.02.24.639789

**Authors:** Mahender Anumalla, Apurva Khanna, Margaret Catolos, Joie Ramos, Ma Teresa Sta. Cruz, Challa Venkateshwarlu, Jaswanth Konijerla, Sharat Kumar Pradhan, Sushanta Kumar Dash, Yater Das, Dhiren Chowdhury, Sanjay Kumar Chetia, Janardan das, Phuleswar Nath, Girija Rani Merugumala, Bidhan Roy, Navin Pradhan, Monoranjan Jana, Indrani Dana, Suman Debnath, Anirban Nath, Suresh Prasad Singh, Khandakar Md Iftekharuddaula, Sharmistha Ghosal, Mohammad Ali, Sakina Khanam, Md Mizan Ul Islam, Muhiuddin Faruquee, Hosna Jannat Tonny, Md Rokebul Hasan, Anisar Rahman, Jauhar Ali, Pallavi Sinha, Vikas Kumar Singh, Mohammad Rafiqul Islam, Sankalp Bhosale, Ajay Kohli, Hans Bhardwaj, Waseem Hussain

## Abstract

Developing high-yielding, flood-tolerant rice varieties is essential for enhancing productivity and livelihoods in flood-prone ecologies. We explored genetic avenues beyond the well-known *SUB1A* gene to improve flood resilience in rice. We screened a collection of 6,274 elite genotypes from IRRI’s germplasm repository for submergence and stagnant flooding tolerance over multiple seasons and years. This rigorous screening identified 89 outstanding elite genotypes, among which thirty-seven exhibited high submergence tolerance, surpassing the survival rate of *SUB1A* introgression genotypes by 40-50%. Thirty-five genotypes showed significant tolerance to stagnant flooding, and 17 demonstrated dual tolerance capabilities, highlighting their adaptability to varying flood conditions. The genotypes identified have a broader genetic diversity and harbor 86 key QTLs and genes related to traits such as grain quality, grain yield, herbicide resistance, and various biotic and abiotic traits, highlighting the richness of the identified elite collection. Besides germplasm, we introduce an innovative breeding approach called ‘Transition from Trait to Environment’ (TTE). TTE leverages a parental pool of high-performing genotypes with complete submergence tolerance to drive population improvement and enable genomic selection in the flood breeding program. Our approach of TTE achieved a remarkable 65% increase in genetic gain for submergence tolerance, with the resulting fixed breeding genotypes demonstrating exceptional performance in flood-prone environments of India and Bangladesh. The elite genotypes identified herein represent invaluable genetic resources for the global rice research community. By adopting the TTE approach, which is trait agonistic, we establish a robust framework for developing more resilient genotypes using advanced breeding tools.

**Plain Language Summary:** To address climate challenges, an urgent focus is necessary to identify and develop flood-tolerant rice varieties, particularly for flood-prone ecosystems across Asia and Africa. We screened 6,274 elite genotypes from IRRI’s germplasm and identified 89 promising lines with improved tolerance to submergence and stagnant flooding. Among these, 37 demonstrated 40-50% greater submergence tolerance than *SUB1A* introgression lines, 35 exhibited stagnant flooding tolerance, and 17 showed dual tolerance. These genotypes contain 86 key QTLs and genes associated with yield, grain quality, and biotic and abiotic tolerance traits. A new breeding strategy, the ‘Transition from Trait to Environment’ (TTE) approach, was developed. We achieved a genetic gain of 65% for submergence tolerance in rice using this method. The newly identified germplasm provides invaluable genetic resources for the global rice research community to develop flood-tolerant rice genotypes.

**Core ideas:**

⍰ The *SUB1A* gene, enabling rice to survive underwater for 14 days, marked a significant breakthrough.
⍰ We have identified elite genotypes with submergence tolerance significantly surpassing the *SUB1A* gene-mediated tolerance.
⍰ The diverse elite genotypes identified harbor 86 key genes and QTLs that affect various traits positively.
⍰ Developed a unique breeding strategy for implementing population improvement in challenging environments.
⍰ The new breeding strategy demonstrated a genetic gain of 65% for submergence tolerance.

## 1 Introduction

Rice (*Oryza sativa* L.) is the staple food for nearly half of the world’s population, with South Asia and Southeast Asia contributing 65% of global rice production. By 2050, the demand for rice in these regions is expected to increase by 87% (Radanielson et al., 2018). The global population is expected to reach 10 billion by 2050 (Yu & Li, 2022; Feeding the future global population, 2024). Much of the population increase will occur in Africa and Asia, regions highly dependent on rice as food (Solis et al., 2020; Rawat et al., 2022). The rice ecologies in Asia and Africa, ranging from lowland to deep-water ecosystems, are negatively impacted by flash floods or prolonged flooding over approximately 20 million hectares in Asia and significant areas in Africa (Bailey-Serres et al., 2010; Dar et al., 2017; Panda & Barik, 2021). Climate change is anticipated to increase the patterns, severity, and frequency of flooding in the coastal regions of Asia and Africa (Hirabayashi et al., 2013; Al-Tamimi et al., 2016; Bailey-Serres et al., 2019; Konapala et al., 2020), posing significant challenges to rice production, especially in lowland coastal regions where it can cause substantial losses (Ismail et al., 2012; Kato et al., 2019). Flood-prone rice ecosystems account for 7% of the global rice area and contribute 4% to global rice production (Yang et al., 2017). The increased demand for rice production and sustainable food security can be met if marginal environments, like flooding ecologies, can dependably provide stable rice yields (Melino & Tester, 2023). There is a heightened need to bridge this gap and a huge opportunity to develop next-generation rice cultivars with broader resilience to fluctuating and prolonged flooding at any stage of crop growth.

In flood-prone ecologies, rice fields experience various flooding conditions. Short-term flash floods or submergence can completely inundate rice plants during the seedling or vegetative stage for a few days to 2 to 3 weeks (Bailey-Serres et al., 2010, 2012; Voesenek & Bailey-Serres, 2013, 2015). Long-term partial submergence, also known as stagnant flooding, involves water depths of approximately 20-50 cm and can persist for weeks to months (Kato et al., 2019; Panda & Barik, 2021; Lin et al., 2024). Deep-water flooding occurs in areas with water levels ranging from 0.5 to 4 meters, lasting for most of the growing season (Catling, 1993; Bailey-Serres & Voesenek, 2008; Hattori et al., 2009). These diverse flooding scenarios pose significant challenges to rice production and require adaptive strategies to mitigate their effect.

Rice plants have developed two strategies to cope with flooding: quiescence and escape (Bailey-Serres et al., 2010, 2012; Voesenek & Bailey-Serres, 2013; Lin et al., 2024). The escape strategy is predominant in deep-water or floating rice ecologies, where vigorous internode elongation keeps the plant’s upper parts above water, facilitating normal gas exchange between submerged and aerial environments (Parlanti et al., 2011). This elongation is driven by the ethylene-responsive factor genes *SNORKEL1 (SK1)* and *SNORKEL2 (SK2),* identified in the deep-water rice variety C9285 (Hattori et al., 2009; Daniel & Hartman, 2024). In contrast, the quiescence strategy is the main adaptive mechanism in flash flooding or submergence, where shoot or leaf elongation is limited. This mechanism is regulated by the ethylene response factor at the *SUB1* locus, which conserves energy reserves and limits shoot elongation to enhance survival and promote regrowth after floodwaters recede (Xu & Mackill, 1996; Xu et al., 2000; Toulotte et al., 2022). The *SUB1* contradicts *SK1* and *SK2* by limiting the shoot elongation and conserving the energy reserves to increase survival and resume growth after the flooding water recedes (Panda & Barik, 2021).

The identification of the submergence tolerance gene *SUB1A* on chromosome 9 in FR13A, an Indian flood-tolerant rice variety derived from the traditional landrace Dhalputtia, marked a significant scientific breakthrough (Xu & Mackill, 1996; Xu et al., 2000). The *SUB1* locus, which includes three ethylene-responsive factor (ERF) genes—*SUB1A, SUB1B*, and *SUB1C*— was pivotal in this discovery (Xu et al., 2000, 2006). Among these, *SUB1A* was confirmed as the key gene conferring submergence tolerance. This breakthrough successfully introduced *SUB1A* into several major rice varieties, enabling them to withstand flash floods for up to two weeks (Septiningsih et al., 2009; Bailey-Serres et al., 2010; Ismail et al., 2013a; Singh et al., 2016; Emerick & Ronald, 2019). One notable variety, Swarna-Sub1, was developed by introgressing the *SUB1A* gene through marker-assisted selection (MAS) and has been celebrated for its potential to aid rice farmers in flood-prone regions of India (Neeraja et al., 2007; Ismail et al., 2013b). Following this success, several submergence-tolerant rice varieties were developed by incorporating *SUB1A* into popular varieties through marker-assisted selection (MAS), enhancing the resilience of rice crops against flooding (Septiningsih et al., 2009; Rumanti et al., 2018; Yamano, 2023).

The *SUB1A* gene is renowned for its excellent submergence tolerance, but its effectiveness diminishes under fluctuating and prolonged submergence of 10 days or more (Singh et al., 2009, 2017; Gonzaga et al., 2016, 2017). The *SUB1A* introgression lines do not match the tolerance level of FR13A, with survival rates dropping to 40-60% under extended and varied flooding conditions (Hussain et al., 2024). Rice researchers have been exploring additional genetic resources to identify new genes and enhance submergence tolerance in rice varieties (Toojinda et al., 2003; Septiningsih et al., 2012; Gonzaga et al., 2016, 2017; Singh et al., 2016; Ismail, 2018; Khalil et al., 2024). However, unlike the *SUB1A* gene, significant success in identifying and leveraging new genes for submergence tolerance is yet to be achieved.

In real-world conditions, stagnant flooding and submergence often occur together, complicating the development of rice varieties that can tolerate both stresses. Despite the co-existence of stagnant flooding and submergence in farmers’ fields, integrating both tolerance levels in the same genotype have not been evident. Overall, progress in genetic improvement to tolerate stagnant flooding and its integration with submergence remains limited (Kato et al., 2019).

In summary, the *SUB1A* gene alone is insufficient to provide complete submergence tolerance for extended periods and under fluctuating flooding conditions, especially in the context of climate change (Shu et al., 2023; Hussain et al., 2024). Progress in integrating submergence and stagnant flooding tolerance into a single genetic background has been limited. Therefore, it is crucial to supplement *SUB1A* with additional tolerance genes and combine it with stagnant flooding tolerance to develop next-generation rice varieties with enhanced resilience to flooding. This work introduces future elite (high yielding, with superior agronomic performance and grain quality) rice germplasm capable of surviving submergence for up to three weeks with over 85% survival rates, significantly outperforming the *SUB1A* gene alone. Additionally, it presents next-generation elite germplasm with high yield and broader resilience to stagnant flooding. In addition to germplasm development, we present a new breeding approach called a transition from trait to environment (TTE) for successfully implementing population improvement and genomic selection for success under challenging environments like flooding. The TTE approach is based on the unique concept of fixing first tolerance to submergence in the parental pool and crossing only those parents tolerant to submergence, subsequently shifting the focus to yield and other agronomic traits under natural flooding conditions. The TTE breeding approach is highly trait-agnostic and can be leveraged for other abiotic stresses like salinity, drought, and heat in rice and other crops.

## 2 Materials and Methods

### 2.1 Experiment 1: Submergence experiments

#### 2.1.1 Plant materials and phenotypic screening

In this study, a collection of diverse 6,274 genotypes were evaluated to identify the most submergence and flooding-tolerant genotypes (Table S1). Out of 6,274 genotypes, a diverse core collection of 1,422 genotypes sourced from various research programs at the International Rice Research Institute (IRRI) in Los Baños, Laguna, Philippines, was included in the study. The remaining genotypes of 4,852 were fixed elite breeding lines, including parental lines from IRRI’s flooding breeding program. The 6,274 genotypes are derived from 888 unique cross-combinations with 89 diverse donors and landraces (Table S2). All 6,274 genotypes used in this study were genotyped using 1k-RiCA SNP markers data (Arbelaez et al., 2019).

The submergence screening was done using IRRI’s standard protocol, with the modification of extending the submergence period to three weeks rather than two weeks (Figure S1). Screening all the rice genotypes for three weeks has been the major intervention in IRRI’s flooding breeding program and the key to identifying the submergence tolerant lines performing beyond the *SUB1A* gene (Anumalla et al., 2023; Hussain et al., 2024). All the submergence field trials were conducted at the IRRI, Los Banos, Philippines, from 2020 to 2023, spanning wet and dry seasons. The dry season (DS) period typically occurs from January to May, and the wet season (WS) is the rainy season, which starts in June and ends in December (Sing et al. and Das et al., 2009). The first experiment began in 2020 DS, and subsequently, only those genotypes were selected for the next screening with a survival rate of more than 85%. After multiple rounds of 4-5 submergence screenings across the seasons and years, the top-performing submergence tolerant genotypes were extracted for grain yield and quality trait evaluation under normal field conditions at IRRI, HQ, Los Baños (Figure S2).

#### 2.1.2 Experimental design

The initial experiments involving large sets of a thousand lines were carried out using an augmented randomized complete block design (RCBD). For subsequent experiments involving a few hundred genotypes selected after the initial screening, we employed an Alpha lattice design with two replications, planting each genotype in three rows of 3 m² length. Twenty-one-day-old seedlings were transplanted at 20 cm x 20 cm spacing, with one seedling per hill across three rows. The tolerant check FR13A was planted along the border and distributed randomly within the experimental field for validation. Additionally, other tolerant checks included Swarna-Sub1, IR64-Sub1, Sambha Mahsuri-Sub1, Ciherang-sub1, IRRI 119, IRRI 224, and IRRI 235. Alongside these, susceptible checks such as Swarna, IR64, Sambha Mahsuri, and IR 42 were also included in the experiments.

#### 2.1.3 Data collection and analysis

In all the submergence experiments, survival percentage data were gathered at 7, 14, and 21 days following the de-submergence or water removal from the field after three weeks. For the data analysis, we consider the 14-day survival percentage most appropriate to classify the genotypes as tolerant or non-tolerant (Hussain et al., 2024). The survival rate of each genotype is calculated by dividing the number of hills that survived post-de-submergence by the total number of hills planted before submergence, then multiplying by one hundred. In this experiment, the average survival percentage across replications served as a criterion to identify the most tolerant genotypes. Genotypes exhibiting over 85% survival were selected for further screening.

### 2.2 Experiment 2: Stagnant flooding

#### 2.2.1 Phenotypic screening and experimental design

We adhered to IRRI’s standard protocol for screening all 6,274 genotypes for stagnant flooding tolerance at IRRI’s experimental fields (Figure S3). In brief, the water depth was increased weekly by 5 cm, reaching 40 cm by 56 days after transplanting (DAT), 50 cm by 63 DAT, and 70 cm by 70 DAT. The tolerant checks included IRRI 119 and Jalmagna, while the susceptible checks were Swarna-Sub1, IR64-Sub1, Sambha Mahsuri-Sub1, IRRI-154, and Swarna. Like submergence experiments, the initial experiments involving large sets of a thousand lines were carried out using an augmented RCBD design. We employed an Alpha lattice design with two replications for subsequent experiments involving a few hundred genotypes selected after the initial screening. Twenty-one-day-old seedlings were transplanted with one plant per hill, spaced 20 × 20 cm apart, using an augmented RCBD design.

#### 2.2.2 Data collections

To better identify elite lines tolerant to stagnant flooding, we expanded our data collection beyond grain yield (kg/ha). Other traits we consider for selecting the best stagnant flooding tolerant genotypes include plant type at maturity (erect and compact), tiller numbers, culm diameter, number of internodes, and internode elongation. We calculated the shoot elongation ratio (SER) to evaluate how quickly or slowly each genotype elongates in flooding conditions. The SER was derived using the following formula:

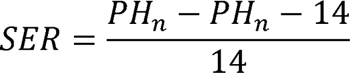

where (n) is the number of days after transplanting, and PH is the plant height (cm) at different water depths during various growth stages. The SER traits evaluated included the shoot elongation rate from the 2nd to 4th weeks (SER_10cm), 4th to 6th weeks (SER_40cm), 6th to 8th weeks (SER_50cm), and 8th to 10th weeks (SER_65cm) after transplanting. For the culm diameter, we measured the central part of the second internode from the plant base in two directions using a vernier caliper. Additionally, data on days to flowering and initial plant height (cm) were collected seven days after transplanting (DAT) while maintaining a water depth of 10 cm. The final plant height was measured during harvesting, with three plants per row in two replicates.

#### 2.2.3 Statistical analysis

A single trial analysis was performed each season for grain yield data (kg/ha) and other agronomic traits described above. Using our data analytical pipeline, mixed models were implemented to analyze and correct the data for the experimental design factors and spatial trends (correlated residuals across rows and columns) (Hussain et al., 2022). Among the five models, the best model was selected based on the lower AIC values and residual plot information to extract the Best Linear Unbiased Predictors (BLUPs) for all the collected traits. Each season, the best genotypes were advanced and screened in subsequent seasons. The baseline model used for running the mixed models is as follows:

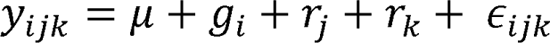

where,

*y_ijk_* is the effect of *i*th genotype in *j*th replication and *k*th block

μ is the overall mean

*g_i_* is the random effect of the *i*th genotype

*r_j_* is the fixed effect of *r*th replication

*r_k_* is the random effect of *k*th block

*∈_ijk_* is the residual error

In the above model, if the design was augmented RCBD, the replication effect was replaced with a block effect, and blocks were used as random.

In the five mixed models, **Model 1 and 2,** we assume residuals are independent and identically distributed as ∈∼*iidN(0, σ^2^_∈_)*. In models 3, 4, and 5, we assume residuals are correlated based on the distance between plots along both the rows and columns, where ∈∼*N(0, R)* and **R** is the covariance matrix of ∈. **Model 3** assumes the structure of the covariance residuals **R** = σ^2^_∈_ Σ_c_ (ρ_c_) ⊕ Σ_r_(ρ_r_), where σ^2^_∈_ is the variance of spatially dependent residual; Σ_c_(ρ_c_) and Σ_r_(ρ_r_) represents the first-order autoregressive correlation matrices and ρ_c_ and ρ_r_ are the autocorrelation parameters for the columns and rows; ⊕ represents the Kronecker product between separable auto-regressive processes of the first order in the row-column dimensions. In **Model 4**, *R= I_C_. σ^2^_∈_ ⊕ Σ_ro_(ρ_ro_)*, where I represents an independently and identically distributed variance structure for columns, and in **Model 5**, *R= I_C_. σ^2^_∈_ ⊕ Σ_c_(ρ_c_) ⊕ I_r_*, where I_r_ represents an independently and identically distributed variance structure for rows.

Comprehensive details of all models and their descriptions are provided in our analytical pipeline and are available on GitHub (https://github.com/whussain2/Analysis-pipeline).

### 2.3 Experiment 3: Yield evaluation of elite panel

#### 2.3.1 Plant materials and phenotypic evaluation

For yield evaluation, we utilized 89 genotypes from a total collection of 6,274 genotypes intended for a future elite panel with better flooding tolerance. Among these 89 genotypes, 35 show high tolerance to stagnant flooding, 37 demonstrate strong submergence tolerance, with over 85% survival rates after three weeks of submergence, and 17 are tolerant of stagnant flooding and submergence. The yield experiments took place in IRRI’s experimental field from 2022, spanning two years and three seasons. We used an Alpha lattice design with two replications. Each genotype was sown in four rows within a 5m² plot. IRRI’s standard fertilizer application and agronomic management protocols were followed throughout the experiments to guarantee consistent and reliable results.

Each genotype’s data on grain yield (kg/ha), days to flowering, maturity, plant height, and grain quality parameters were observed. The data were collected according to IRRI’s standard procedures. In addition to these traits, data on grain quality parameters, such as grain length, width, amylose content, head rice recovery, chalkiness, and gelatinization temperature, were estimated for all the genotypes.

#### 2.3.2 Phenotypic data analysis

For this experiment, multi-environment analysis was performed across two years and three seasons among the identified 89 genotypes using grain yield and other agronomic traits. Multi-environment trial (MET) analysis was performed using a single stage-wise approach (Hussain et al., 2022). The mixed model fitted is given below in the matrix notation:

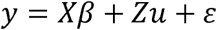

Where,

*y* is a m x 1 vector of individual phenotypes

*X* is a m x n design matrix relating phenotypes to fixed effects of replications and trials

β is a vector of fixed effects

*Z* is a design matrix assigning phenotypic individuals to the marker effects

*u* is a random effect of genotypes

ε is n x n matrix of residual/error effects

Here we assume u has variance=*Var(u)∼a^2^_g_G*, where *G* is genomic or kinship co-variance matrix of n x m dimensions (n is no. of markers and m is no. of individuals) representing genomic similarity of individuals, and var(ε)= σ^2^_e_*I, I* is identity matrix.

The genomic relationship matrix () was constructed using the equation:

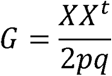

Here, ***X*** is a scaled and centered matrix of marker data, and p and q are the frequencies of the major and minor alleles in marker data. In the above model, genomic estimated breeding values (GEBVs) were obtained to characterize the 89 genotypes.

#### 2.3.3 Genotypic characterization and diversity analysis

The best 89 genotypes were accessed for genetic diversity and the presence of harboring significant abiotic and biotic tolerant QTLs/genes using 1k-RiCA genotypic data. For genetic diversity analysis, 1k-RiCA genotypic data was used to construct the genomic relationship matrix (GRM). Principal component analysis was done on GRM to visualize the diversity and relationship among genotypes as a biplot. Distance-based on the ward’s method (Strauss & Maltitz, 2017) was used to construct the distance matrix and build a dendrogram.

The diversity of the 89 genotypes in relation to the 3K genome (Wang et al., 2018) was also assessed. For this, the marker coordinates based on 1k-RiCA genotypic data were extracted from the 3K genome marker data file available in the Rice SNP-seek Database (https://snp-seek.irri.org/). Using markers with similar coordinates, the genotypes from the 1k-RiCA and 3K genomes were combined, and a new genomic relationship matrix (GRM) was constructed. PCA was done on the GRM and visualized as a biplot to compare the diversity and relationship of 89 genotypes with the 3K genome Indica group. The description of the 89 genotypes, with detailed information, is provided in Table S3.

All the analyses were performed using R software (R Core Team 2023). The mixed models were fitted using the *ASReml-R* package (Butler et al., 2018). The genetic diversity analysis was conducted in R using *FactoMineR* and *Factoextra* R packages (Lê et al., 2008). The genomic relationship matrix was built using the R package *AGHMatrix* (Amadeu et al., 2016). Diversity among the genotypes was visualized using the biplot created using the R package. The variables for the biplot were obtained through the principal component analysis (PCA) performed on GRM using the function *princomp* in R software. Genetic gains were estimated by regressing the mean survival scores for submergence tolerance on the year of evaluation of the genotypes as per the procedure detailed in (Khanna et al., 2022, 2024).

## 3 Results

### 3.1 Phenotypic features of the elite pool

After rigorous multi-year and multi-season testing, the best 89 genotypes out of 6,274 diverse collections were identified as elite breeding resources for flooding tolerance. The elite pool was subdivided into three groups: 37 submergence-tolerant, 35 stagnant flooding-tolerant, and 17 tolerant to both submergence and stagnant flooding. We classified genotypes as submergence tolerant if underwater for 21 days of submergence there was an average survival score of more than 85%. Stagnant flooding tolerance (low to moderate tolerance to submergence) was classified by an average survival score of less than 50%, and stagnant and submergence tolerant genotypes with a more than 60% survival score for submergence tolerance (Table S3). We categorize the genotypes into three groups because submergence tolerance and stagnant flooding tolerance have antagonistic mechanisms (Kato et al., 2019). It is essential to have germplasm resources that are tolerant to submergence, stagnant flooding, or even both.

### 3.2 Submergence-tolerant elite genotypes

The 37 submergence elite lines identified in this study were the best submergence-tolerant genotypes, as their survival percentages were significantly more than the comparator *SUB1A* introgression genotypes (Figure 1A). Eight *SUB1A* introgression genotypes are currently being cultivated in the farmer’s fields in Asia as flooding-tolerant genotypes were the comparators (Ismail et al., 2013a). Significant differences in survival rate were observed among the genotypes and in the interaction between season and year (Table S4). The differences across the years and seasons were non-significant, indicating the stability in the survival score among the identified genotypes. The mean survival percentage of the newly identified elite genotypes was 86.52 ± 10 (Figure 1B). On the other hand, the mean survival percentage of the *SUB1A* introgression genotypes was significantly lower at 33.39 ± 16. The survival score of the *SUB1A* genotypes was even lower than those we classified as stagnant flooding-tolerant genotypes. The survival score of the submergence-tolerant genotypes was closer to the FR13A, a donor for the *SUB1A* gene, which is globally recognized as one of the best genotypes tolerant to submergence (Hussain et al., 2024).

**Figure 1.**
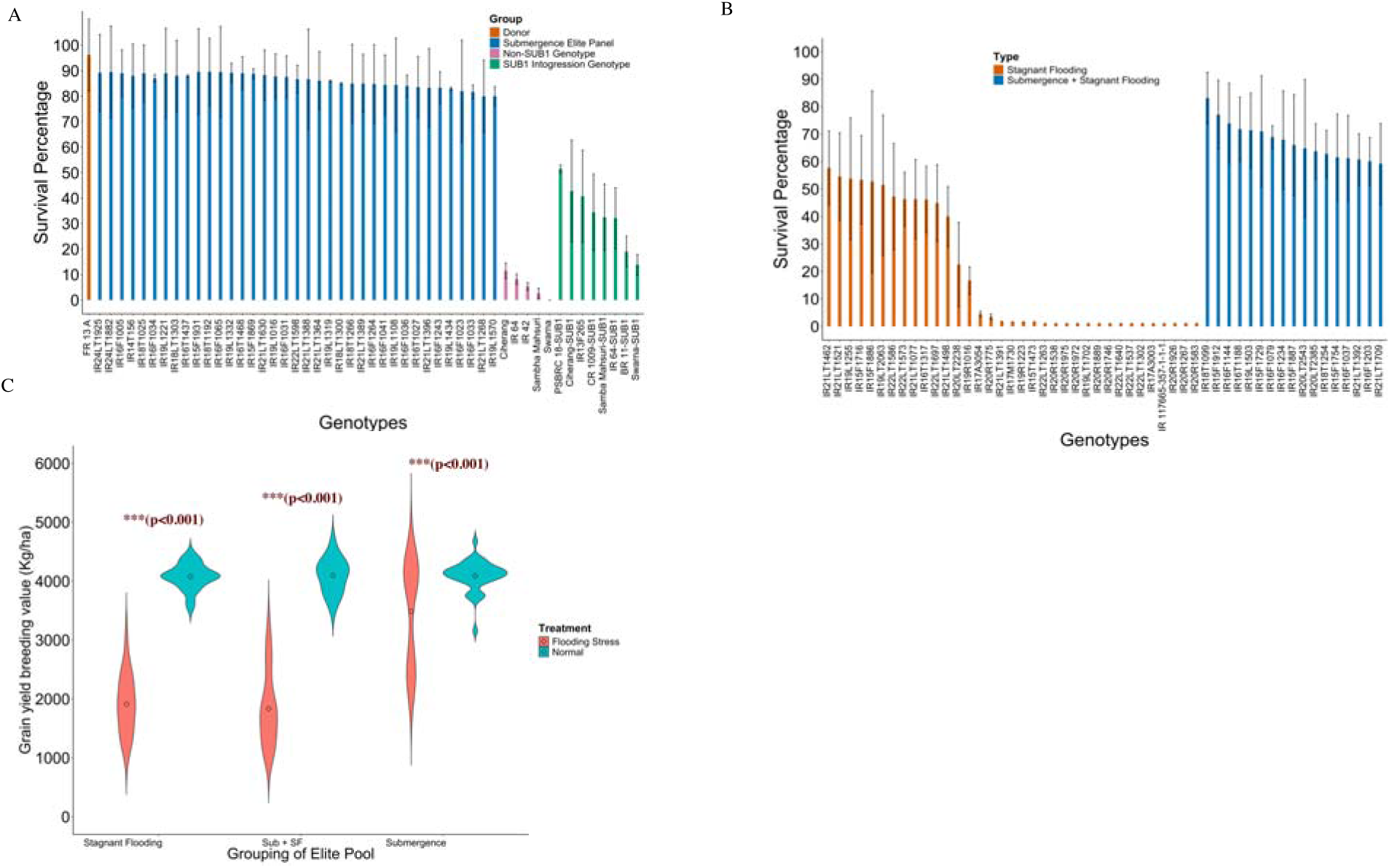
Illustrates the morphological features and characteristics of the elite pool identified in this study. (A) Displays the submergence tolerance elite genotypes identified in this research. The data is averaged across 5-6 experiments, with the bars representing the standard errors for each genotype. The y-axis indicates the survival percentage, while the x-axis lists the names of the elite genotypes. The elite genotypes shown (in blue bars) demonstrate a tolerance level to submergence that is equivalent to that of donor FR13A. The eight *SUB1A* introgression genotypes exhibit significantly lower submergence tolerance compared to the identified elite genotypes. (B) Depicts the submergence tolerance of genotypes with high stagnant flooding tolerance. We categorized these into two types based on survival percentage. Genotypes with more than 60% survival are classified as both stagnant flooding and submergence tolerant genotypes (blue bars), whereas genotypes with less than 60% survival are classified as only stagnant flooding tolerant (orange bars). (C) Presents the breeding value for grain yield (kg/ha) for three groups.

In addition to the submergence tolerance, we chose genetically diverse genotypes with high grain yield and superior agronomic performance. The breeding value for grain yield of the submergence tolerant genotypes under normal conditions ranges from 3500 to 5500 (kg/ha), with a mean of 4093.96 ± 129 (Figure 1C). The genotypes we identified had head rice recovery (HRR) ranging from 43.50% to 70.5% (Table S3).

Plant height ranged from 96.68 to 134.39 (cm), and days to maturity from 114 to 142 days. The grains of the new pool of elite genotypes identified had different shapes, sizes, and amylose content, including short, medium, and long slender grain types. The amylose content ranged from intermediate to high, with values of 17.50 to 28.10 (Table S3).

### 3.3 Stagnant flooding-tolerant elite genotypes

Only 35 genotypes were identified as the best stagnant flooding-tolerant genotypes from a collection of 6,274 genotypes. The criteria for choosing the best stagnant flooding tolerant genotypes were high grain yield, a tolerant ideotype of erect flag leaf, compact plant type, strong stem and diameter, and greater internode elongation ability (Figure 1B, Figure S5). Besides these traits, we also considered the grain quality performance, the presence of essential genes for abiotic and abiotic stresses, and genetic diversity among the genotypes. The breeding value for yield under normal conditions varied from 2589.55 (kg/ha) to 4498.18 (kg/ha), with a mean breeding value of 4032± 545 (kg/ha). In the stagnant flooding conditions, the breeding values for grain yield were lower, ranging from 1055.32 (kg/ha) to 3181.57 (kg/ha) with a mean of 1936 ± 650 (kg/ha) (Figure 1C).

The range of variability in grain yield was more evident in stress conditions than in normal conditions. Besides grain yield, significant plant height and flowering differences were observed under regular and flooding conditions (Table S5, Figure S4). Higher height is expected under stagnant flooding as the plant tends to elongate under stagnant flooding conditions. The stem elongation rate at different depths, tiller numbers, and claim diameters for each genotype are given in Table S5. The genotypes identified had different shapes and sizes and amylose content, including short, medium, and long slender grain types (Table S3). The genotypes identified had high HRR ranging from 41.98 to 70.30%. The amylose content of the selected genotypes ranged from low to high, with values of 14.50 to 26.30%. In addition to tolerance to stagnant flooding, most of the genotypes extracted also showed higher submergence tolerance of 10 to 60% (Figure 1B). The survival rate of 15 genotypes was much higher than that of the *SUB1A* introgression genotypes, thus having higher submergence tolerance and a very high tolerance to stagnant flooding.

### 3.4 Submergence plus stagnant flooding-tolerant genotypes

Of the 6,274 genotypes in the diverse collections, only 17 were identified as having both stagnant flooding and submergence tolerance. We classified the genotypes as tolerable to both stresses if they exhibited a suitable ideotype for stagnant flooding and related features and had more than 65% survival score (Figure 1B). The survival score of the genotypes ranged from 59.19% to 83.15%, with a mean survival score of 69±13%. The grain yield of the genotypes under normal conditions ranged from 3513.01 (kg/ha) to 4672.55 (kg/ha) with a mean of 4093.96 ± 298.11 (kg/ha). Under stagnant flooding stress conditions, the grain yield varies from 1238.75 (kg/ha) to 3181 (kg/ha), with a mean of 1833±634.20 (kg/ha). The difference in plant height and flowering was significant under stress and normal conditions (Table S5, Figure S4). The genotypes identified had different shapes, sizes, and amylose content, including short, medium, and long slender grain types. The genotypes identified had an HRR ranging from 50.90 to 65.50%, amylose content ranging from 14.9 to 26%, and divergent grain shapes and dimensions.

### 3.5 Genetic diversity of identified elite genotypes

A genome relationship matrix was used to assess the genetic diversity of the selected genotypes. The selected 89 genotypes of the elite pool represented substantial diversity, as seen from the dendrogram analysis of the marker data (Figure 2A). The entire pool was clustered into three distinct groups. Each main cluster was further divided into sub-clusters. The arrangement of genotypes in relation to submergence tolerance, stagnant flooding tolerance, or both was random. Therefore, similar genotypes did not necessarily group in similar tolerance types (Figure 2A). The donor for the *SUB1A* gene, FR13A, was part of a small cluster comprising only five genotypes. This suggested that the selected tolerant genotypes came from different genetic backgrounds and could be useful as a future elite breeding resource. This was borne out by pedigree analysis, which demonstrated that the lines originated from varied parental and donor backgrounds. Originally, a total of 67 diverse founder lines were used to generate the entire collection of 6,274 genotypes (Table S2 and Table S6). Among these, 20 founder lines were frequently used in breeding programs with different genetic backgrounds (Table S6).

**Figure 2.**
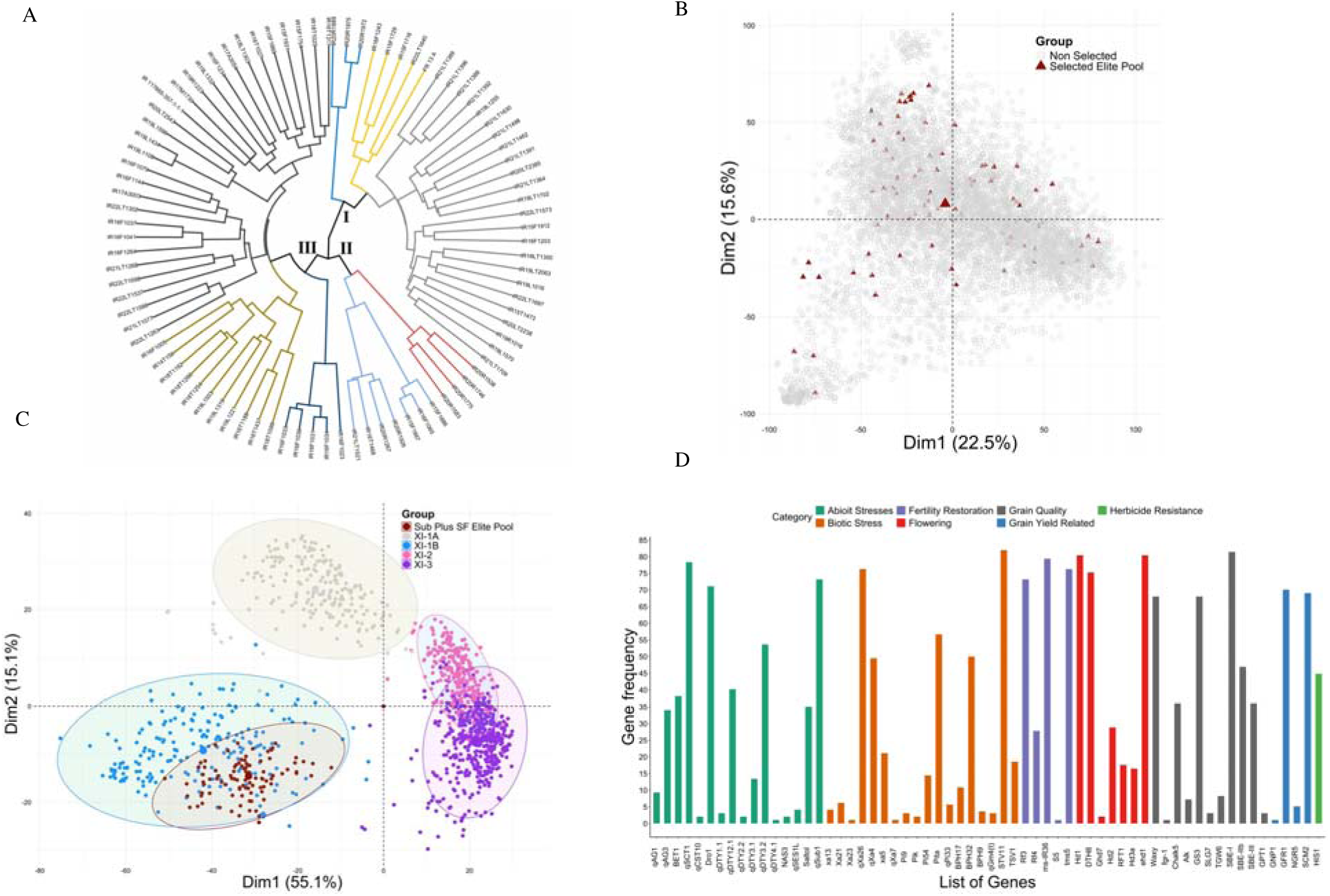
Genetic characterization and diversity analysis of the identified elite pool of 89 genotypes. (A) shows the dendrogram of the 89 elite genotypes based on the distance using the Wards method by leveraging SNP marker data. The genotypes were grouped into three main groups, and each group was subdivided into subgroups, indicating the diverse nature of genotypes. The FR13A donor for submergence tolerance falls in small group, indicating the diverse nature of other genotypes for flooding tolerance. (B) shows the representation of the identified elite pool of 89 genotypes (dark red triangles) from the whole collection of 6,274 genotypes. (C) shows the representation of the identified elite pool among the 3K genome Indica group. The elite lines identified fall in the *XI-1B* group, representing the modern breeding cultivars bred by IRRI and in Asia. (D) Illustrates the frequency of major genes in the identified elite pool of 89 genotypes, with the x-axis showing a list of present genes and the y-axis indicating the frequency of these genes. The genes are organized and color-coded based on various trait categories.

The biplot of principal component analysis (PCA) derived from the genomic relationship matrix illustrated the diversity of the selected lines and their representation of the entire collection of 6,274 genotypes (Figure 2B). The random and dispersed nature of the identified 89 genotypes among the whole collection indicates the representation of the entire diversity of the whole collection present in the identified elite collection of 89 genotypes.

The genetic and functional diversity of rice is exemplified by 3,000 (3K) genome accessions from 89 countries (Li et al., 2014). We utilized genotype data from the 3K genome panel to gain insights into the diversity of our identified elite pool and how well it represents the diversity of the 3K genomes, specifically within the Indica group (Wang et al., 2018). Historically, rice has been grouped into the Xian/*Indica* (XI)and *Japonica* groups. The *Indica* group is divided into XI-1A from East Asia, XI-1B comprising modern varieties from various origins, XI-2 from South Asia, and XI-3 from Southeast Asia. Notably, all elite pool lines clustered in the XI-1B group (Figure 2C), which represented the modern breeding cultivars primarily bred at IRRI and in Asia (Xie et al., 2015).

### 3.6 High-value genes and QTLs in the identified genotypes

Since the selected pool derived primarily from elite breeding lines developed at IRRI and other countries in Asia, an analysis for genes/QTLs for other useful traits, above and beyond tolerance to submergence and stagnant flooding, revealed the presence of several biotic and abiotic stress-related genes/QTLs. Genes related to abiotic stresses like drought and cold and other biotic stresses such as bacterial blight and blast were identified. Additional genes were related to desirable alleles for fertility restoration, flowering, grain quality, grain yield, and herbicide resistance (Figure 2D). The detailed list of 86 genes found in the elite pool associated with different categories and traits and their frequencies are given in Table S7. For example, the frequency of the *SUB1A* gene in the elite pool is 73%. This also suggested that higher tolerance achieved than that with *SUB1A* would be due to additional genes and synergistic mechanisms. Other critical genes of importance included salinity tolerance genes with *Saltol* QTL at a frequency of 35%. Besides these, the elite pool harbors six QTLs for drought tolerance, two additional QTLs for salinity tolerance, eight genes for bacterial blight, nine genes for blast resistance, three genes for BPH tolerance, five genes for fertility restoration, two genes for anaerobic germination, three genes for cold tolerance, one gene each for Fe toxicity, boron toxicity tolerance, gall midge, stripe virus, and tungro virus resistance, amylose content, fragrance, chalkiness, tillering number, and herbicide tolerance. The details of genes and associated trait category information are provided in Table S8.

### 3.7 Utility of the new elite pool

After the identification of the elite tolerant genotypes, in 2020, we selected the top 10 genotypes with a submergence survival rate above 85% from our elite collection and made 30 crosses. By the 2024 dry season, we had 627 fixed genotypes (stage 1, F_6_ derived) from these crosses, which we evaluated for submergence tolerance by submerging them for 21 days. More than 50% of genotypes showed excellent tolerance, with survival rates over 75% (Figure 3A, Table S9). The average survival rate for the entire population, including susceptible checks and eight *SUB1A* introgression genotypes, was around 68% (Figure 3A). Comparing these results with previous stage 1 trials, we observed a significant improvement in survival rates, demonstrating the superior performance of the genotypes selected in 2020 for submergence tolerance.

**Figure 3.**
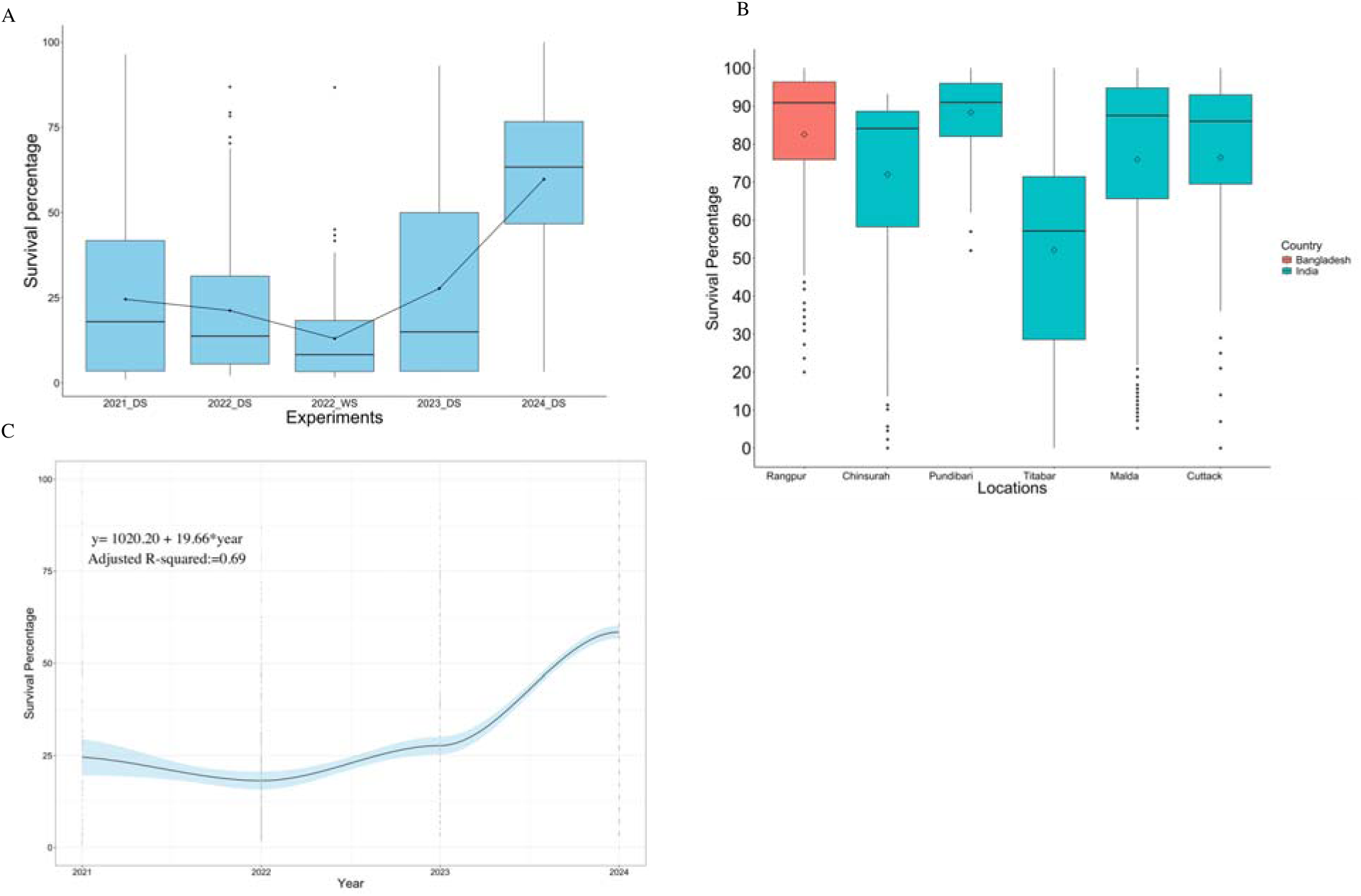
Shows the usefulness and importance of the elite pool in generating the next-generation flood tolerance genotypes. (A) Comparison of the fixed breeding lines across the years for submergence tolerance. Each year represents a different set of materials generated in IRRI’s submergence breeding program by crossing different parents yearly. The 2024 stage 1 or fixed genotypes represent the materials generated using the parents with submergence tolerance fixed based on TTE breeding approach. The significantly higher survival percentage compared to other stage 1 trials shows the usefulness of the new breeding approach and the materials for flooding tolerance. (B) shows the survival percentage of the cross materials across diverse flooding ecologies of India and Bangladesh. Again, the high survival percentage of genotypes in the natural flooding ecologies demonstrates the usefulness of the new materials generated with the TTE approach. (C) Genetic gain for submergence tolerance using the last 3 years of data from the stage 1 trial. The big jump in 2024 in survival percentage and increase of 65% shows the potential and usefulness of the elite x elite crossing based on the TTE breeding approach.

In addition to assessing the performance of these materials at IRRI-HQ, we collaborated with the National Agriculture Research Extension Service (NARES) partners in India and Bangladesh to evaluate these genotypes across ten natural flooding ecologies in 2024. Of these ten locations, only five in India and one in Bangladesh experienced flooding for 10-15 days. The genotypes demonstrated high submergence tolerance across all locations (Figure 3B), with an average survival rate of 90%, except at Titabar, where the average survival rate was 50%. The most tolerant genotypes were common and had survival rates exceeding 80%. We also analyzed genetic gains by regressing survival scores on the year trials were conducted at IRRI-HQ for submergence tolerance. This revealed a higher gain in 2024, with an overall genetic gain of 65% (Figure 3C). These findings highlight a significant success in enhancing submergence tolerance using our newly identified elite genotypes and the uniquely designed TTE breeding approach, detailed in the discussion section

## 4. Discussion

Despite the discovery of new genes and QTLs for submergence and stagnant flooding tolerance (Sripongpangkul et al., 2000; Gonzaga et al., 2016; Singh et al., 2016; Ismail, 2018), the development of commercial rice varieties with comprehensive flooding tolerance remains elusive. This is particularly true for varieties that can endure both stagnant flooding and submergence beyond the capabilities provided by the *SUB1A* gene. The urgency for rice genotypes with enhanced flood resilience has never been greater. Farmers are increasingly confronted with unpredictable flooding, exacerbated by climate change (Hirabayashi et al., 2013; Hauer et al., 2020), which can impact crops at any growth stage. Identifying germplasm capable of withstanding such conditions is crucial.

Our research on future flooding-tolerant rice elite germplasm represents a significant leap forward in rice flood breeding efforts. These elite genotypes exhibit high tolerance to flooding and possess high grain yield, superior agronomic performance, and genetic diversity. These elite genotypes serve as reservoirs of key genes and QTLs related to various agronomic, grain quality, biotic, and abiotic traits, making them invaluable genetic resources for population improvement-based breeding strategies.

### 4.1 Cultivate resilience beyond *SUB1A*

The *SUB1A* gene has set a benchmark for submergence tolerance in rice during short-term flooding events (Bailey-Serres et al., 2010). Significant strides have been made in incorporating the *SUB1A* gene into popular varieties cultivated across millions of hectares in Asia (Das et al., 2009; Ismail et al., 2013a). However, its effectiveness diminishes under prolonged and fluctuating flooding conditions (Hussain et al., 2024). Our earlier study revealed that submergence lasting beyond 10 days significantly reduced survival rates among the *SUB1A* introgression genotypes (Hussain et al., 2024). Additionally, these genotypes exhibited significant fluctuations in submergence tolerance across different years and seasons, rendering them highly unstable for consistent submergence tolerance (Hussain et al., 2024).

In contrast, the newly identified tolerant elite genotypes demonstrate significantly better submergence tolerance compared to the *SUB1A* introgression genotypes. Their submergence tolerance is 40-50% greater (Figure 1A), and they exhibit higher stability (Table S3). Screening these genotypes for submergence with 21 days underwater was aimed at enhancing their stability and adaptability. For instance, a genotype that survives 21 days of submergence with an 85% survival rate is likely to maintain similar or higher survival rates under 14 days or less across various locations. These selected genotypes also display a broader range of flooding tolerance, combining resilience to both submergence and stagnant flooding.

This dual-stress tolerance is particularly crucial in real-world scenarios where flooding stresses can occur at any stage of crop growth. The 17 genotypes capable of tolerating both types of flooding highlight the potential for integrating dual-stress tolerance mechanisms, thereby reducing the risks and unpredictability associated with climate change impacts in rice cultivation regions. This advancement underscores the importance of developing rice varieties with enhanced flood resilience to ensure stable and sustainable rice production.

### 4.2 Readily available valuable genetic resources

Rice landraces and other accessions in the gene bank hold immense potential for identifying genotypes with broad resilience to flooding conditions. However, directly utilizing these genotypes in breeding programs has been challenging due to linkage drag and the time-consuming process (Toulotte et al., 2022). Additionally, efforts to identify and integrate additional genes beyond *SUB1* have faced significant hurdles. Our prolonged submergence screening for 21 days and targeted selection based on phenotypes have successfully captured genetic variation for *SUB1A* and additional polygenic variation for submergence tolerance. The elite pool also includes a substantial genetic variation for stagnant flooding tolerance, making both stagnant flooding and short-term submergence-tolerant genotypes valuable and readily available genetic resources.

In addition to resilience to submergence and various flooding stages, the identified genotypes have been carefully selected for high agronomic performance, including grain yield and grain quality. The genotypes identified vary in growth duration, amylose content, and grain type to meet global demands and preferences. For instance, in flood-prone areas of Bangladesh, both medium and late-duration germplasm with high amylose content is required (Iftekharuddaula et al., 2019), while in India, flood-prone areas mainly need long-duration germplasm with intermediate amylose content.

The elite genotypes are valuable sources of biotic and abiotic tolerance QTLs and genes. The elite pool harbors genes and QTLs for various stresses, including drought, cold, salinity, bacterial blight, and blast (Figure 2D, Table S7). The *SUB1A* gene is present in most of the elite pool genotypes and is completely fixed. Additionally, the elite pool possesses genes for traits such as fertility restoration, flowering, grain quality, grain yield, and herbicide resistance. Thus, the developed elite pool is not only a source of flood tolerance but also a reservoir of key genes and QTLs crucial for breeding future climate-resilient rice varieties.

The elite genotypes identified are derived from 67 founders and through various breeding schemes ranging from single to complex crosses (Table S6), representing the breadth of IRRI’s flooding tolerance breeding program diversity (Figure 2). The substantial genetic diversity within the elite pool is a standout feature that is crucial for any private or public breeding program. This diversity provides a robust genetic base for developing future rice varieties. To ensure elite pool genotypes are genetically diverse and representative of decades of IRRI’s flooding breeding program and varietal development, genomic and pedigree data were used to account for genetic similarity among the genotypes and identify these genetically diverse elite genotypes. The dendrogram and biplot clearly illustrate how the elite genotypes are genetically diverse and represent the diversity of the entire breeding collection (Figure 2A, 2B).

### 4.3 Potential for higher genetic gains

Achieving genetic gains has been a major goal in public and private breeding programs. A set of high-performing elite genotypes with a broader tolerance to flooding is highly required to unlock the potential of cultivation in flooding ecologies and boost genetic gains. The primary motivation for identifying an elite pool of genotypes with superior agronomic performance and tolerance to submergence is to achieve higher genetic gains by leveraging population improvement and genomic selection in the breeding program.

Recurrent selection, a breeding strategy centered on population improvement, has been pivotal in boosting genetic gains (Khadr & Frey, 1965). This efficient scheme aims to increase the frequency of favorable alleles for a given quantitative trait through repeated cycles of crossing and selection (Orf, 2008). Each cycle of crossing and phenotyping results in recombining and reshuffling alleles into the best combinations, which are then selected as parents for the next cycle. This process improves the population’s mean and boosts genetic gains for quantitative traits like grain yield (Rutkoski, 2019). To extract the best candidates from a given cycle and develop better genotypes for varietal release, it is essential to cross the best with the best or elite x elite parents (van Ginkel & Ortiz, 2018). When leveraged with recurrent selection, genomic selection (GS) can help in selecting the best and most reliable genotypes based on genomic estimated breeding values (GEBVs), increasing selection accuracy and accelerating breeding cycles for enhanced genetic gains.

However, traditional flooding rice breeding programs or any breeding programs for abiotic stress-related traits face challenges in effectively implementing population improvement and genomic selection. For example, traditional breeding programs at IRRI have mainly focused on crossing non-elite (donors/landraces) with high-yielding elite breeding lines to develop a high-yielding elite tolerant genotype (Khanna et al., 2022, 2024). The focus has also been on introducing pyramiding-tolerant QTLs for a given abiotic trait in elite backgrounds. However, this approach is inappropriate when the goal is enhancing genetic gains through a population improvement-based breeding strategy (Khanna et al., 2022). First, in this breeding approach, only a few high-yielding tolerant genotypes are identified, which is insufficient to drive recurrent selection due to narrow diversity (Figure 4A). For population improvement, an optimal number of high-performing genotypes is needed to cross, select, and recycle to achieve short-term and long-term genetic gains (Allier et al., 2019; Juma et al., 2021). Second, screening genotypes in real stress-prone environments to identify high-yielding tolerant genotypes results in high variation and standard deviation, leading to low heritability and poor covariance structure among genotypes within and across trials (Barreto et al., 2024; da Costa et al., 2024). This is mainly because genetic variation for tolerance to abiotic stress traits will segregate in the derived breeding population, resulting in high, moderate, and low tolerance genotypes. Stress primarily impacts the growth and development of moderate and low-tolerance genotypes, resulting in poor data on agronomic traits, including yield and high variations within the breeding population. In genomic selection, prediction accuracy is closely related to heritability, which depends on the population’s genetic covariances and error variance structures (Krishnappa et al., 2021).

**Figure 4.**
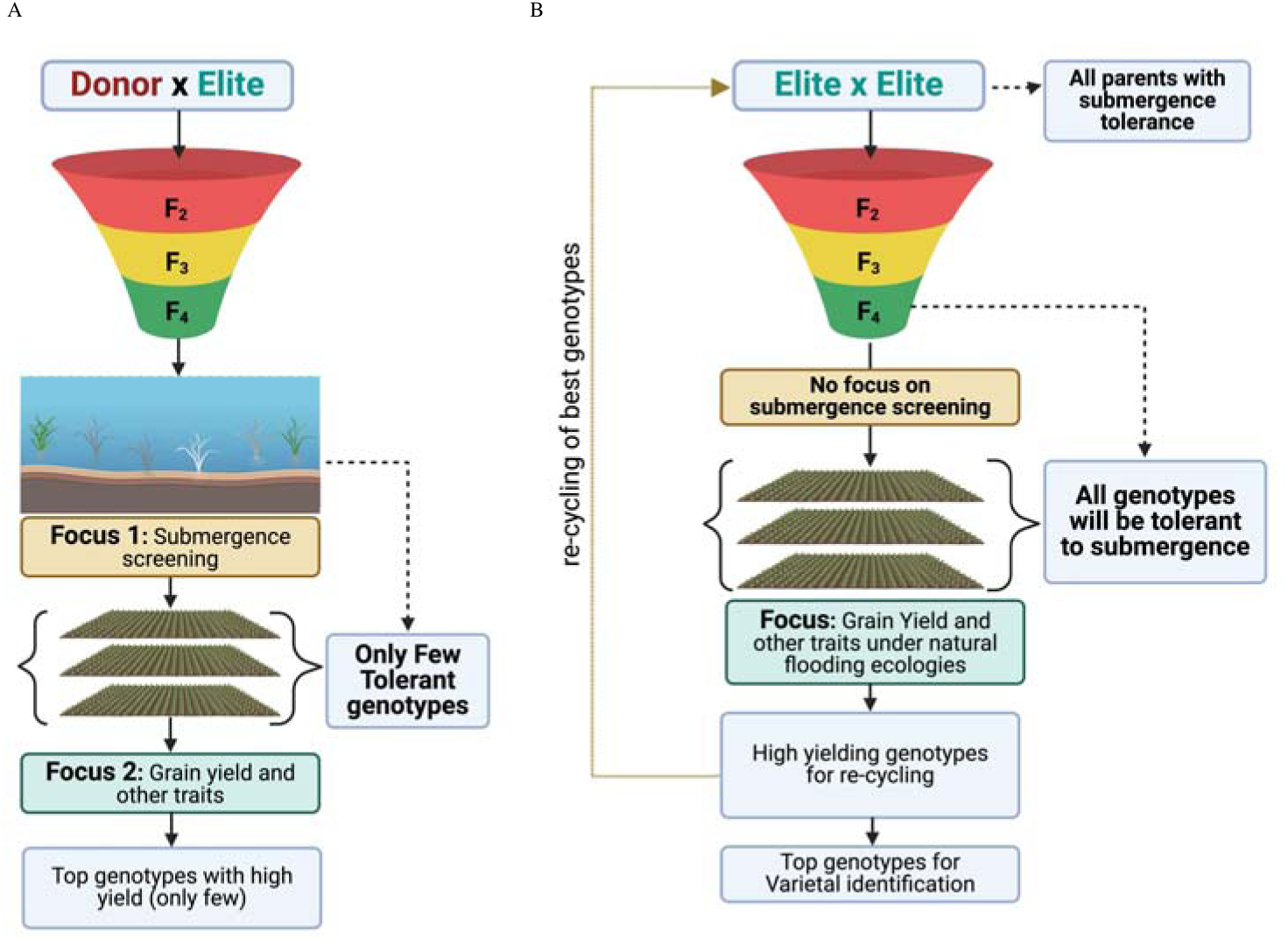
Illustration of the breeding schemes used to develop flood-tolerant genotypes. In (A), the traditional method involves crossing a donor or submergence-tolerant genotype with high-yielding elite genotypes to identify a genotype that is both tolerant and high-yielding. Initially, the focus is screening for submergence tolerance, followed by identifying high-yielding genotypes. However, the likelihood of finding genotypes that exhibit both traits is low, making this approach impractical for population improvement or leveraging genomic selection. In (B), we present our redesigned flood breeding approach and the unique concept of TTE. This method starts with identifying an elite pool of genotypes entirely tolerant to submergence. These genotypes are crossed, and their progenies are evaluated in natural flooding conditions for yield performance. All genotypes are expected to survive, providing optimal yield data. By leveraging genomic selection, we can estimate the breeding values of all genotypes, select those with high breeding values, and recycle them for the next crossing cycle, thereby driving population improvement and genetic gains.

To improve heritability and minimize error variance due to differential responses of genotypes in stress environments, we developed a novel approach called Transition from Trait to Environment (TTE). We hypothesized that fixing submergence/flood tolerance in the parental pool and crossing genotypes with both parents tolerant to submergence would allow us to obtain optimal data on yield and other key agronomic traits for every genotype in the breeding population under evaluation, as all genotypes will be submergence tolerant. Thus, this approach treats submergence tolerance as an environmental condition rather than a trait, as it is already fixed in the parents or the elite pool. This shift allows us to concentrate on enhancing genetic gains for yield and other important agronomic traits (Figure 4B). This approach also enables us to select high-yielding genotypes using genomic estimated breeding values through the genomic selection approach. For instance, if we have 30 to 40 high-yielding genotypes, all submergence tolerant, we can cross these genotypes, ensuring that any progeny derived from the crossing will be submergence tolerant. This eliminates the need for submergence tolerance screening, allowing us to focus on grain yield and other agronomic traits in natural flooding ecologies. By leveraging the grain yield data through genomic selection, we can select the top high-yielding genotypes and recycle them as parents for the next crossing block, thereby efficiently driving the population improvement breeding strategy and boosting genetic gains (Figure 4B).

We have demonstrated the effectiveness of this approach by crossing elite parents with submergence tolerance fixed in both parents. The population derived from the crossing scheme with all parent’s tolerance to submergence exhibited a higher mean survival percentage than previous breeding populations derived using a traditional donor x elite crossing scheme (Figure 3A). More than 50% of genotypes showed high submergence tolerance with a survival percentage greater than 75% (Table S9). A remarkable genetic gain of 65% in just one breeding cycle underscores the potential of this approach in driving genetic gains in rice flooding tolerance breeding programs and generating the next generation of genotypes with broader resilience to flooding. Testing the derived genotypes generated using our new approach across different hotspot flooding locations in India and Bangladesh ensures the real-world applicability of this approach. It exemplifies the effectiveness of TTE in driving population improvement breeding strategy in IRRI’s rice flooding breeding program.

## CONCLUSION

By advancing beyond the *SUB1A* gene, we have pioneered next-generation rice germplasm with exceptional tolerance to both submergence and stagnant flooding. The identified elite germplasm is a valuable, ready-to-use genetic resource for breeding programs and research initiatives worldwide. Researchers can harness these genotypes to develop locally adapted, flood-tolerant rice varieties, delve deeper into the genetic mechanisms of flood tolerance, and uncover new genes. The diversity and resilience of these genotypes ensure they are well-equipped to meet global demands and withstand the challenges posed by climate change. Looking ahead, our ambition is to provide world-class germplasm to our National Agriculture Research Extension System (NARES) partners, setting a benchmark for rice breeding and genetic gains through a population improvement-based strategy integrated with modern tools like genomic selection. The elite lines identified in this study open new avenues for innovation and collaboration, paving the way for a resilient and sustainable rice production system capable of thriving in flood-prone environments. Dissecting the molecular mechanisms underlying the improved tolerance would contribute to a better understanding of the synergistic action of the gene involved.

## Abbreviations

1k-RiCA: 1k-Rice custom amplicon
AIC: Akaike information criterion
BLUPs: Best linear unbiased predictors
ERF: Ethylene-responsive factor
GEBVs: Genomic estimated breeding values
GRM: Genomic relationship matrix
HRR: Head rice recovery
PCA: Principal component analysis
QTL: Quantitative trait loci
RCBD: Randomized complete block design
SER: Shoot elongation ratio
SNPs: Single nucleotide polymorphisms
TTE: Transition from Trait to Environment.

## AUTHOR CONTRIBUTIONS

**Waseem Hussain:** Conceptualization, Data curation, Formal analysis, Investigation, Supervision, Validation, Writing - original draft, Writing - review & editing; **Mahender Anumalla:** Conceptualization, Data curation, Formal analysis, Methodology, Visualization, Writing - original draft, Writing - review & editing; **Apurva Khanna:** Data curation, Formal analysis, Methodology, Writing - review & editing; **Margaret Catolos:** Data curation, Formal analysis, Writing - review & editing; **Joie Ramos:** Data curation, Formal analysis, Investigation, Methodology, Writing - review & editing; **Ma Teresa Sta. Cruz:** Data curation, Formal analysis, Investigation, Methodology, Writing - review & editing; **Challa Venkateshwarlu:** Data curation, Investigation,Writing - review & editing; **Jaswanth Konijerla:** Data curation, Investigation,Writing - review & editing; **Sharat Kumar Pradhan:** Data curation, Investigation, Methodology, Visualization, Writing - review & editing; **Sushanta Kumar Dash:** Data curation, Investigation, Methodology, Visualization, Writing - review & editing; **Yater Das:** Data curation, Investigation, Writing - review & editing; **Dhiren Chowdhury:** Data curation, Investigation, Writing - review & editing; **Sanjay Kumar Chetia:** Data curation, Investigation, Writing - review & editing; **Janardan das:** Data curation, Investigation, Writing - review & editing; **Phuleswar Nath:** Data curation, Investigation, Writing - review & editing; **Girija Rani Merugumala:** Data curation, Investigation, Writing - review & editing; **Bidhan Roy:** Data curation, Investigation, Writing - review & editing; **Navin Pradhan:** Data curation, Investigation, Writing - review & editing; **Monoranjan Jana:** Data curation, Investigation, Writing - review & editing; **Indrani Dana:** Data curation, Investigation, Writing - review & editing; **Suman Debnath:** Data curation, Investigation, Writing - review & editing; **Anirban Nath:** Data curation, Investigation, Writing - review & editing; **Suresh Prasad Singh:** Data curation, Investigation, Writing - review & editing; **Khandakar Md Iftekharuddaula:** Data curation, Investigation, Supervision, Writing - review & editing; **Sharmistha Ghosal:** Data curation, Investigation, Writing - review & editing; **Mohammad Ali:** Data curation, Investigation, Writing - review & editing; **Sakina Khanam:** Data curation, Investigation, Writing - review & editing; **Md Mizan Ul Islam:** Data curation, Investigation, Writing - review & editing; **Muhiuddin Faruquee:** Data curation, Investigation, Writing - review & editing; **Hosna Jannat Tonny:** Data curation, Writing - review & editing; **Md Rokebul Hasan:** Data curation, Investigation, Writing - review & editing; **Anisar Rahman:** Data curation, Investigation, Writing - review & editing; **Jauhar Ali:** Visualization, Writing - review & editing; **Pallavi Sinha:** Project administration, Writing - review & editing; **Vikas Kumar Singh:** Project administration Resources, Writing - review & editing; **Mohammad Rafiqul Islam:** Project administration Resources, Writing - review & editing; **Sankalp Bhosale:** Funding acquisition, Project administration, Resources, Supervision, Writing - review & editing; **Ajay Kohli:** Project administration, Supervision, Validation, Writing - review & editing; **Hans Bhardwaj:** Funding acquisition, Resources, Supervision, Writing - review & editing

## Acknowledgments

The authors thank research technicians Cesar Castillo, Alma Fuentes and Evangeline Angeles for their help in conducting the trials. The authors also want to thank the Breeding Operations Unit (BOU), Enterprise Breeding System (EBS), and other shared services of IRRI for providing all the support and help in generating the data. The authors thank all the IRRI breeders and researchers who have generated the genotypes used in this study and shared seeds with us. Authors also thank NARES research technicians and field workers for support in conducting the trials. The work was primarily supported by the Bill and Melinda Gates Foundation (BMGF) under the Accelerating the Genetic Gains in Rice (AGGRi) project. Funding was also received from the Accelerated Breeding Initiative (ABI) work package under the OneCGIAR.

## DATA AVAILABILITY STATEMENT

All data supporting the findings of this study can be found in the Supplementary Information Tables S1-S9.

## SUPPLEMENTARY INFORMATION

**Table S1:** Details and a list of a diverse collection of 6,274 used in this study.

**Table S2:** Pedigree information and unique parentage of 6,274 genotypes.

**Table S3:** Agronomic characteristics of the identified elite pool of 89 genotypes.

**Table S4:** ANOVA table for the survival scores of elite genotypes screened over multiple years and seasons.

**Table S5:** Phenotypic trait information of 89 genotypes under normal and stagnant flooding stress conditions.

**Table S6:** List of unique founder genotypes used to generate 6,274 genotypes.

**Table S7:** Promising QTLs/gene profile of 89 elite genotypes.

**Table S8:** Detailed description of QTLs/gene information

**Table S9:** Survival rates of Stage 1 genotypes derived from crossing submergence-tolerant parents.

**Figure S1:** Diagram illustrating the 3-week submergence protocol adopted in this study.

**Figure S2:** A schematic representation of the phenotypic screening of the whole collection of 6,274 genotypes.

**Figure S3:** Diagram illustrating the stagnant flooding screening protocol adopted in this study

**Figure S4:** Distribution of days to maturity and plant height (cm) among the 89 elite genotypes under flooding and normal conditions.

**Figure S5:** Visual representation of the tolerant stagnant flooding genotypes grown in the IRRI-HQ field during the 2023 DS

